# CT Multi-Task Learning with a Large Image-Text (LIT) Model

**DOI:** 10.1101/2023.04.06.535859

**Authors:** Chuang Niu, Ge Wang

## Abstract

Large language models (LLM) not only empower multiple language tasks but also serve as a general interface across different spaces. Up to now, it has not been demonstrated yet how to effectively translate the successes of LLMs in the computer vision field to the medical imaging field which involves high-dimensional and multi-modal medical images. In this paper, we report a feasibility study of building a multi-task CT large image-text (LIT) model for lung cancer diagnosis by combining an LLM and a large image model (LIM). Specifically, the LLM and LIM are used as encoders to perceive multi-modal information under task-specific text prompts, which synergizes multi-source information and task-specific and patient-specific priors for optimized diagnostic performance. The key components of our LIT model and associated techniques are evaluated with an emphasis on 3D lung CT analysis. Our initial results show that the LIT model performs multiple medical tasks well, including lung segmentation, lung nodule detection, and lung cancer classification. Active efforts are in progress to develop large image-language models for superior medical imaging in diverse applications and optimal patient outcomes.

## 1 Introduction

The artificial intelligence (AI) paradigm is evolving to train large foundation models on interrelated big datasets via self-supervised learning and adapt the foundation models to empower downstream tasks via transfer learning [BHA^+^21]. Along this direction, large language models (LLM) have achieved great successes, such as GPTs [RNS^+^, RWC^+^19, BMR^+^20], which are used to construct homogenized models and induce emergent functionalities (such as in-context learning). Beyond the LLM, the AI community is actively building multi-modal and multi-task models toward general artificial intelligence (AGI), such as CLIP [RKH^+^21], Gato [RZP^+^22], KOSMOS [HDW^+^23], and GPT-4 [Ope23]. Building a single model across multiple modalities and diverse tasks is promising to improve the performance of individual tasks as it can synergize multiple information sources for a given task.

Inspired by the rapid advancement of the AI field, researchers in the medical imaging field are actively developing or adapting modern AI methods in various medical applications such as medical tomographic imaging [Wan16, WYDM20, WBJ^+^ 22]. Over the past years, important progresses in self-supervised learning, large language models, visual-language models, and so on provide great potential to revolutionize medical imaging.

Self-supervised learning is a key mode to train large models using unannotated big data [NZWL20, NLF^+^22, NSW22, NXW]. Among various self-supervised learning methods [GSK18, CBJD18, NZWL20, ZJM^+^21, NXW], contrastive learning [CKNH20, RKH^+^21], masked modeling [DCLT18, HCX^+^22], and generative modeling [RNS^+^, RWC+19, BMR^+^20] are currently the three most popular that have been successfully applied to large models due to their simplicity and efficiency. In medical image analysis, self-supervised pretraining methods have been explored [ZSP^+^21, TYL^+^22, NW22]. For example, Models Genesis [ZSP^+^21] was proposed to pretrain neural networks by restoring 2D/3D patches from different transformations, demonstrating that the pretrained models can lead to an improved performance of classification and segmentation tasks. Swin UNETER [TYL^+^ 22] was preptrained for image segmentation by minimizing the combination of three self-supervised learning losses, involving contrastive learning, inpainting, and rotation. Based on a generic 3D transformer, URCTrans [NW22] was pretrained via contrastive learning, demonstrating its effectiveness for lung nodule detection. However, to our best knowledge, there is no unified large model up to now that works for multiple medical tasks using tomographic scans by integrating various training datasets with diverse annotations.

The visual-language or image-text model is a most important multi-modal model, such as CLIP [RKH^+^21] and ViLBERT [LBPL19], which combines visual and textual information to perform various visual-language tasks. For example, in the medical domain visual-language models have been studied on analysis of chest X-ray radiographs and associated radiology reports [ZJM^+^22, ZCZ^+^ 22, HPLY23, BHL^+^23]. In particular, ConVIRT [ZJM^+^22] was proposed to learn chest X-ray image representations via bidirectional contrastive learning with image-text pairs, which dramatically reduces the need for labels in downstream classification tasks. REFERS [ZCZ^+^22] was proposed as a cross-supervised method to train a vision transformer on multiview radiographs using free supervision signals from the original radiology reports, which benefited downstream tasks. Furthermore, the X-VL model [HPLY23] was designed to explore a combination of masked modeling and contrastive learning strategies to pretrain vision and text encoders, which achieved promising results in five downstream tasks. Beyond the alignment of a single image and the corresponding report, BioViL-T [BHL^+^23] was intended to account for prior images and reports when available during both training and fine-tuning stages. Nevertheless, there has been no visual-language model that works on 3D medical images and the associated reports. The development of large visual-language models for 3D tomographic scans is an exciting research opportunity for major reasons; for example, low-dose chest CT scans can provide much more detailed information and are routinely used for lung cancer screening.

Currently, LLMs (e.g., ChatGPT) are mainly used as a tool or an API for different purposes, such as patient discharge summary [PL23], medical application writing [Bis23], radiology decision-making [RKK^+^23], cardiovascular disease prevention recommendations [SBVI^+^23], clinical report translation [JSD^+^22, LTZ^+^23], clinical report de-identification [LYZ^+^23], and ensemble learning on Chest X-ray images [WZO^+^ 23]. However, it has not been demonstrated so far how to synergize current LLMs and LIMs in the cases of large 3D image models for improved diagnostic performance.

Based on the above analysis, we are motivated to develop a multi-modal multi-task model leveraging LLM and LIM models. Our large image-text (LIT) model is designed in a unified framework and can be trained using a wide range of interrelated medical datasets with diverse annotations. We envision such unified models can maximally extract the information hidden in interrelated big datasets and effectively incorporate comprehensive prior knowledge, ultimately forming a versatile intelligent agent to aid physicians and deliver the best possible diagnosis, treatment, and prognosis of various diseases. In this study, we primarily focus on multi-task learning related to clinical report generation from chest CT scans. High-dimensional tomographic images present at least two unique challenges to the development of large visual-language models relative to the cases of 2D images such as radiographs. First, processing high-dimensional 3D chest CT images is computationally demanding, making the use of large models rather challenging. Second, super-sparse annotations (for example, only a small lung nodule is reported) in clinical reports make the training of image captioning models highly non-trivial. To overcome the first challenge, we leverage cost-effective training techniques. For the second challenge, instead of directly generating clinical reports, our large model decomposes the clinical report into multiple tasks first, performs these multiple tasks simultaneously, and then generates the clinical report. In this stage for feasibility demonstration, we mainly focus on semantic segmentation, object detection, and classification tasks using 3D chest CT images. The initial results have demonstrated the feasibility of our LIT model, which is designed to accommodate additional tasks and data modalities in the future.

The rest of this paper is organized as follows. In the next section, we describe our methodology. In the third section, we report our representative results. In the last section, we discuss relevant issues and conclude the paper.

## 2 Methodology

### 2.1 Overall architecture

The basic idea is to design a unified LIT model trainable on multi-modal medical datasets with various annotations and functional in multiple tasks by synergizing both patient-specific and task-specific prior knowledge through LLM. Currently, we focus on image and text inputs but the idea can be naturally extended to more modalities and more tasks. As shown in Figure 1, our model consists of four components: an image encoder, a text encoder, a task attention module/block, and task decoders. Specifically, the image and text encoders extract image and text features. The task attention layers extract task-specific features from image features and text features conditioned on a task token. The task token specifies task-specific prior and informs which task will be performed, which can be automatically learned or generated by the text encoder. Finally, specific task decoders are used for prediction. In the following, we will describe each of these components in detail.

**Figure 1:**
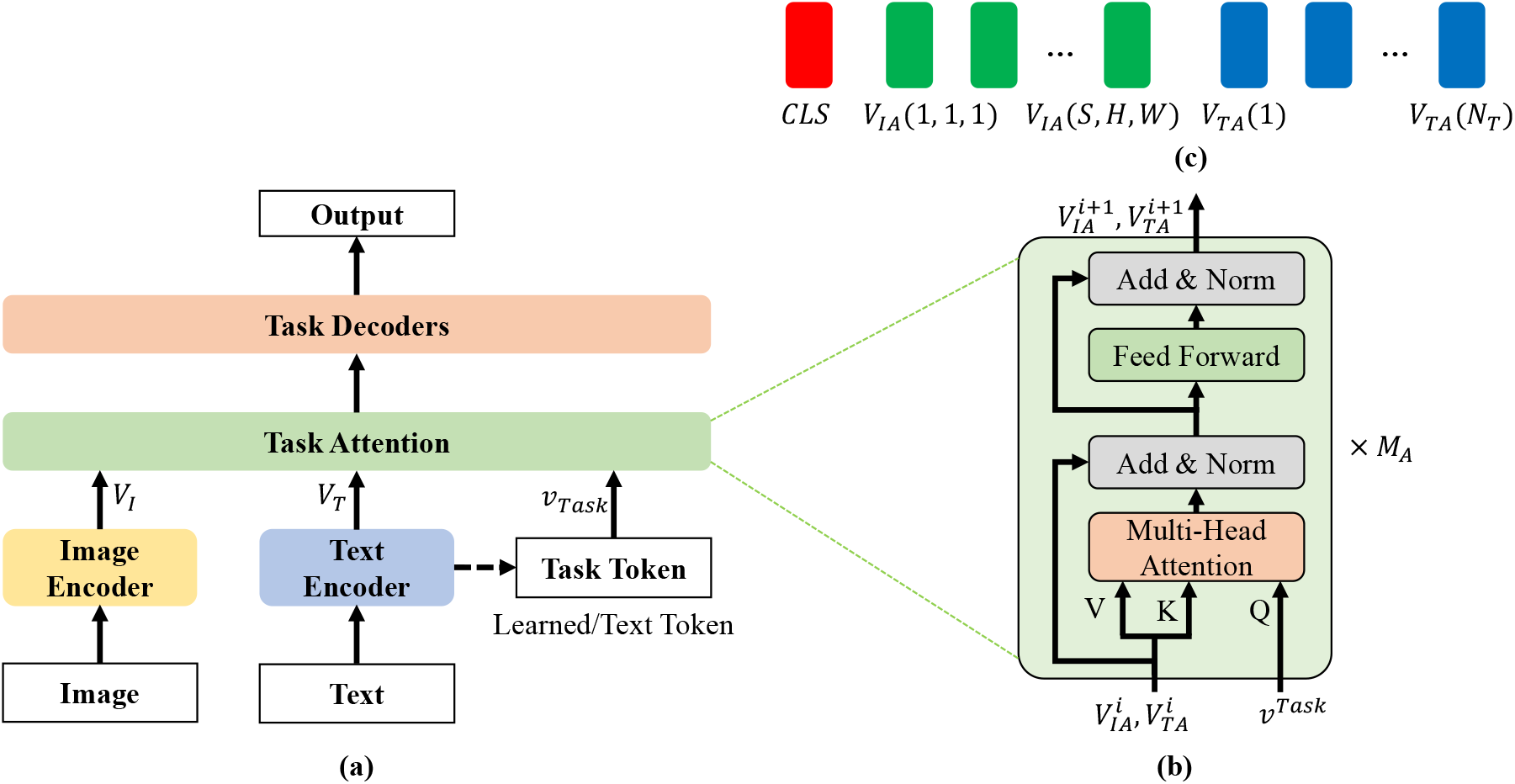
Large image-text (LIT) model for multiple medical CT tasks. (a) The overall architecture of the LIT model, (b) the task attention module, and (c) its output tokens.

### 2.2 Image Encoder

In our LIT model, the plain ViT [DBK^+^20] is implemented as the image encoder. To process a 3D CT scan, we divide each image volume into cubes as in [NW22] and use a linear layer to map the cubes to a sequence of visual tokens. Different from [NW22], here we learn the positional embedding automatically and decompose it into two parts indexing in-plane and through-plane positions respectively. In other words, we have two positional embeddings, one for the 2D space within each slice and the other for the 1D range of slice position. The volumetric positional embeddings are the sum of them. By doing so, the number of learned parameters can be reduced.

Let us denote the size of a 3D CT scan by *S* × *H* × *W*, where *S* is the number of slices and varies significantly from case to case due to different imaging protocols, and *H* and *W* are the dimensions of a slice, and usually *H* = *W* = 512. To keep the input information complete, it is natural is to use the original slice dimensions, i.e., *H* = *W* = 512, without any cropping and down-sampling, and set *S* to proper values in different tasks. However, the full slice resolution means a large 3D input size; *e.g.*, 32 × 512 × 512 is significantly larger than the commonly used sizes, such as 96 × 96 × 96. Here the window attention mechanism [VSP^+^17] is adapted to process full-resolution slices. Specifically, this mechanism spatially divides cube features into non-overlapping windows and restricts the attention to the assigned window. Similar to [FLH^+^22], transformer blocks are evenly divided into four parts, the last block implements global attention to propagate information across windows while the other blocks use window attention. By doing so, the computational cost can be significantly reduced.

### 2.3 Text Encoder

Any existing large language model can be utilized as the text encoder in our proposed model. Here we explored the use of the language model developed in the biomedical NLP domain, i.e., BioGPT [LSX^+^22], and in the natural visual-language domain, i.e., CLIP [RKH^+^21], as our text encoder. On one hand, the text encoder can encode patient-specific prior in the free text format; e.g., “The patient is 65 years old, a male, the race is white, and has a smoking history of 94.0 pack years.” On the other hand, the text encoder can encode task-specific information as well; e.g., “Classification of adenocarcinomas”, or encode more prior information, such as “Classification of adenocarcinomas, which appear as nodules with solid opacity, nodules exhibiting some ground-glass opacity (GGO) with some solid components (part-solid nodules), or nodules exhibiting complete GGO (ground-glass nodules).” [KY20]. In this way, prior knowledge can be embedded into a unified fashion as prompts to optimize the model prediction.

### 2.4 Task Attention

To perform multiple tasks, we design a task attention component as shown in Figure 1-(b). This task attention extracts task-specific features given a task token. Specifically, the task attention component consists of multiple modified transformer blocks, where the only difference from the original transformer lies in the way of calculating self-attention; *i.e.*, the queries for all elements in all transformer blocks are calculated with the same task token instead of the input tokens themselves. In the task attention component, the task-specific self-attention is computed as

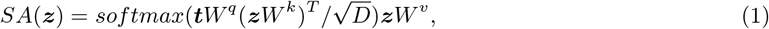

where ***z*** ∈ *R^N×D^* is the input sequence, *N* is the sequence length, *D* is the embedding dimension, ***t*** ∈ *R*^1×*D_t_*^ is the task token, *D_t_* is the task token dimension, *W^t^* ∈ *R^D_t_×D^, W^k^* and *W^ν^* ∈ *R^D×D^* are the two linear mapping matrices respectively. Clearly, the task-token-related signals in all elements are extracted from multi-modal data. In practice, task tokens can be automatically learned or generated by the text encoder. The output sequence of the task attention component includes three parts: CLS token, visual features, and textual features, as shown in Figure 1-(c), which are forwarded to the task decoders accordingly.

### 2.5 Task Decoders

Different tasks are performed with dedicated decoders respectively given task-specific features. Compatibility, scalability, and effectiveness are the main considerations in designing task-specific decoders. All the decoders only take the features from the last layer of the task attention block as inputs, as recent progress has shown that predictions from the last transformer layer are even better than the counterparts using multi-layer features [LMGH22]. This strategy also benefits compatibility and scalability for multiple task-specific decoders. As shown in Figure 2, we have three decoders for classification, segmentation, and detection tasks respectively.

**Figure 2:**
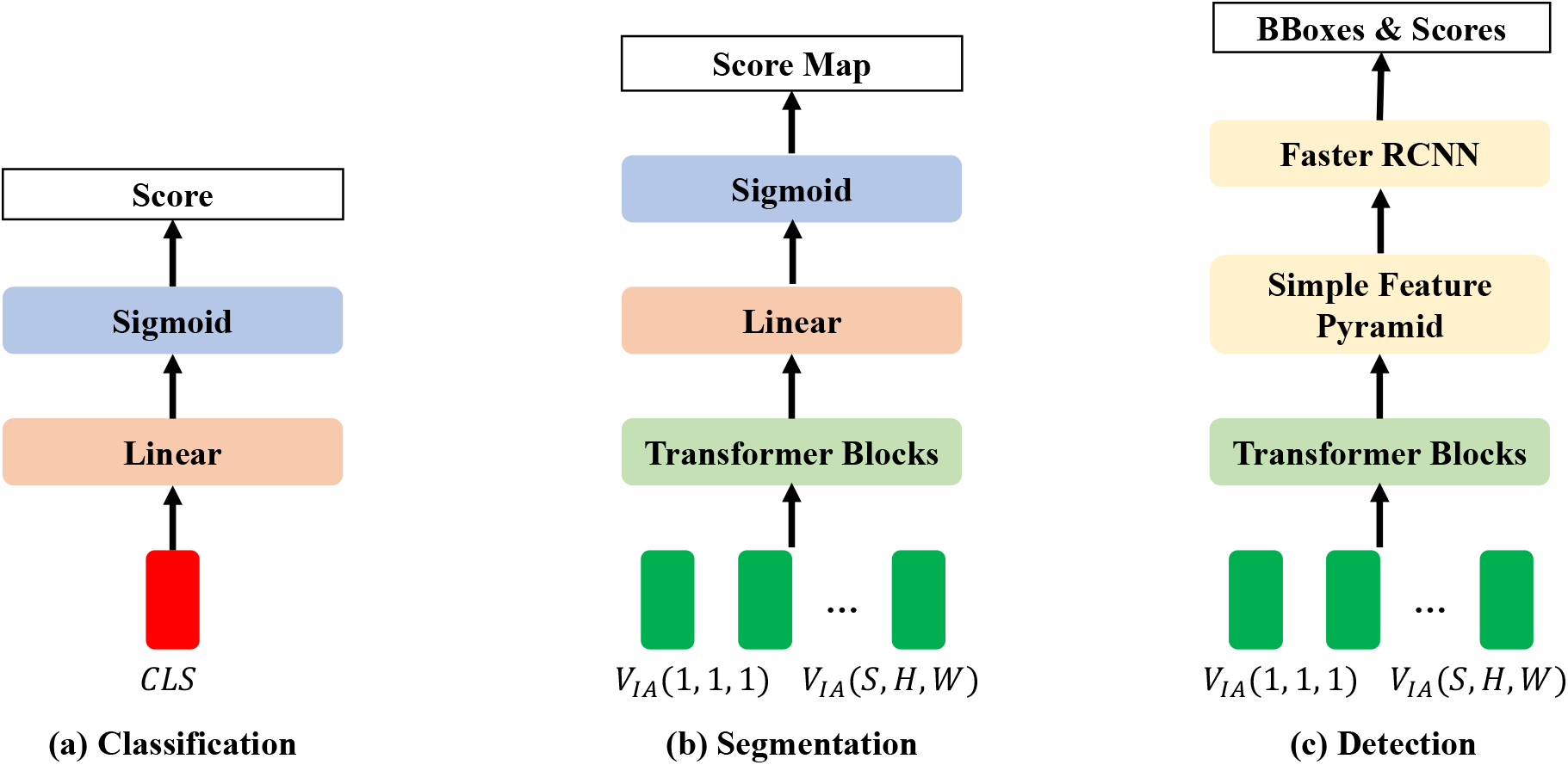
Task-specific decoders for (a) classification, (b) semantic segmentation, and (c) object detection respectively.

For classification, a simple binary classifier is implemented as a linear layer followed by a sigmoid function, taking the CLS token as the input.

For segmentation, the transformer blocks are first used to extract segmentation-specific features, which are then mapped to the original image space using a linear layer. By doing so, the segmentation score map can be directly computed without using any transposed convolution blocks or upsampling operations as previously used [ZLZ^+^20]. This linear mapping can be regarded as a transposed operation upon embedding of image tokens. Here the inputs are the corresponding image features.

For detection, the transformer blocks are used to extract detection-specific features. Then, ViTDet [LMGH22] is adapted for detection. Specifically, VitDet calculates a feature pyramid from single-layer features, where different feature maps are used to detect objects/structures of different sizes using the Faster RCNN detector [RHGS15]. Note that various other object detectors can be implemented as needed.

By combining our task attention blocks with task-specific decoders, all tasks can be performed in a prompting mode so that multi-modal data, patient-specific, and task-specific priors can be aggregated through the self-attention mechanism toward optimal prediction. In our implementation, we tend to set the number of decoder parameters as small as possible to maximize the use of the powerful representation ability of the pretrained large models.

### 2.6 Self-Supervised Pretraining

To optimize large models, self-supervised pretraining at scale is the key step. In our model, the image encoder and text encoder were pretrained on big image and text datasets. There are three main approaches for pretraining language models; *i.e.,* masked modeling of BERT, generative modeling of GPT, and contrastive learning. Since the pretraining strategies of large language models have been extensively studied, we respectively used the pretrained CLIP in the natural visual-language domain and the pretrained BioGPT in the biomedical NLP domain as our text encoder.

For pretraining vision models, contrastive learning and masked autoencoding are two commonly used selfsupervised learning methods. Here we adapted the masked autoencoder method [HCX^+^22, FLH^+^22] to pretrain our image encoder on 3D CT datasets. Briefly speaking, the image encoder introduced in Subsection 2.2 was optimized by predicting masked cubes (90%) from a small number of visible cubes (10%), using an additional shallow transformer decoder. To reduce the memory overhead, only a part of the slices along the longitude direction were predicted while recovering each cube.

### 2.7 Multi-Task Training

After self-supervised pretraining, our multi-task model in Figure 1-(a) can be trained either separately or simultaneously, which is in contrast to most of the current studies that only finetuned different models for different tasks. As discussed before, unified training can leverage interrelated datasets with diverse annotations to optimize our large model. On the other hand, the proposed model will simplify the deployment without storing multiple model weights. Since a prompting mode is used to achieve multi-task, we correspondingly arranged the training data in a prompt style. Taking the segmentation of left and right lungs as an example, a sample was first randomly selected, then a prompt label was randomly selected, and its prompt text was constructed; *e.g.,* “Segmentation of left lung”, and finally the binary mask of either the left or right lung was randomly selected. The mask values consistent with the prompt label were set to 1, and the other mask values set to 0. This training data sampling strategy was used for all other tasks. Currently, we fixed our text encoder and optimized the parameters of the rest of the components in the multi-task training stage. The binary cross-entropy loss function was used in the classification and segmentation tasks. The losses used for the object detection task were the same as those in [LMGH22], and only a single class of the lung nodule was used for the detection task.

## 3 Experimental Design and Results

### 3.1 Tasks and Datasets

In this feasibility study, we focus on lung cancer diagnosis-related tasks, including left/right lung segmentation, lung nodule detection, and lung cancer classification with an emphasis on identifying adenocarcinoma from small cell carcinoma, large cell carcinoma, and squamous cell carcinoma. We pretrained the image encoder using 124,731 3D CT scans selected from the NLST dataset^1^, where each scan with more than 64 slices was selected. The LUNA16 dataset [STdB^+^16] was used for left/right lung segmentation and lung nodule detection tasks. The LUNG-PET-CT-Dx^2^ dataset was used for the lung cancer classification task. For each task, we randomly constructed a training dataset and a testing dataset. The specific configurations are shown in Table 1.

**Table 1:** Configuration of task-specific training and testing datasets.

| Task-specific Data | Number of Training Samples | Number of Testing Samples | Sample Size |
| --- | --- | --- | --- |
| Segmentation | 800 | 88 | $16 \times 512 \times 512$ |
| Detection | 2,003 | 633 | $16 \times 512 \times 512$ |
| Classification | 12,211 | 5,847 | $4 \times 128 \times 128$ |

### 3.2 Results

The encouraging results are shown in Figure 3, demonstrating that our unified model is feasible to accomplish multiple tasks simultaneously. As shown in Table 1, our model can take different sizes of images as inputs, which means that it can be also extended to other modalities, such as 2D Chest X-ray images and 4D CT scans. Our initial quantitative results on representative datasets are reported in Table 2, where our model performance measures were compared using different text encoders, showing that pretrained BioGPT using biomedical data achieved significantly better results than the CLIP counterpart in distinguishing multiple lung cancer types under the same settings. Since the lung segmentation task is relatively easy and only a single task token was used in object detection, different text encoders did not produce any significant difference.

**Figure 3:**
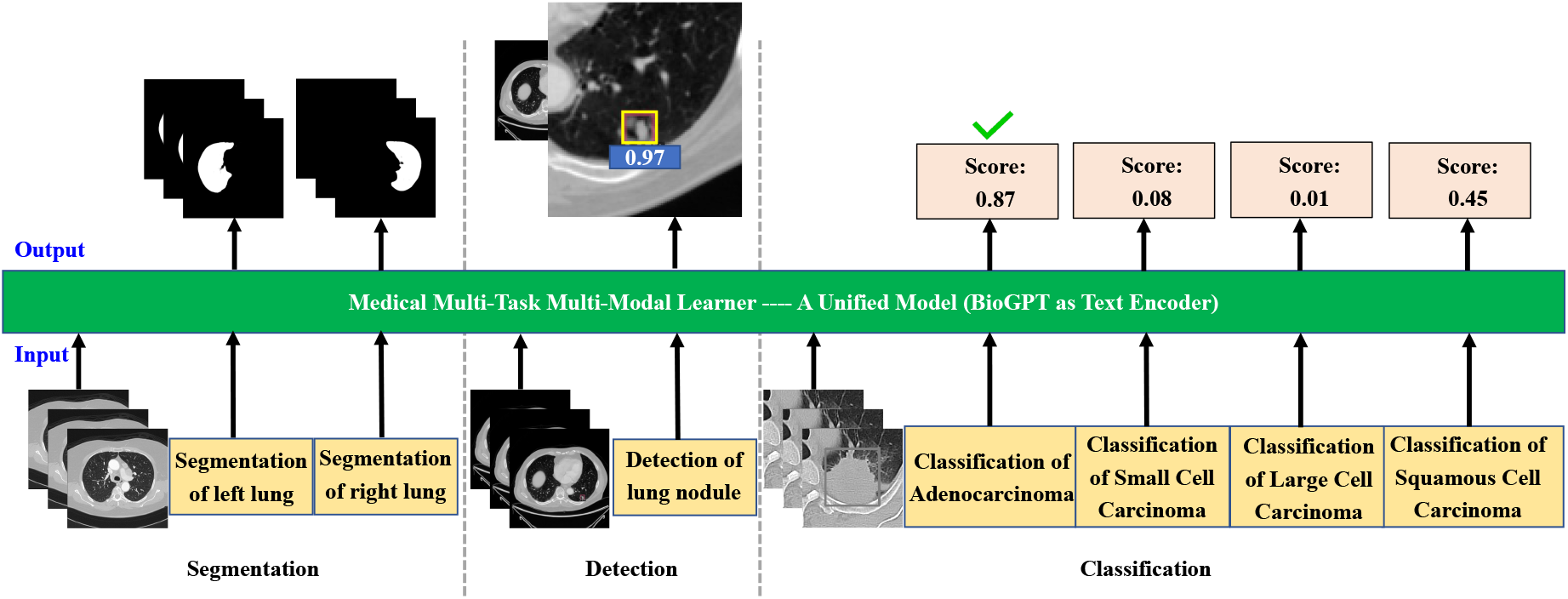
Task-specific decoders for (a) classification, (b) semantic segmentation, and (c) object detection respectively.

**Table 2:** Quantitative results obtained using our proposed LIT model on typical datasets using CLIP and BioGPT, respectively. Intersection-over-Union (IoU), average precision (AP), and accuracy (ACC) values are reported for segmentation, detection, and classification tasks respectively.

| Model | Segmentation (IoU in %) | Detection (AP in %) | Classification (ACC in %) |
| --- | --- | --- | --- |
| LIT-CLIP | 95.8 (left:96.1/right:95.5) | 50.3 | 79.5 |
| LIT-BioGPT | 95.9 (left:96.0/right:95.8) | 50.3 | 85.1 |

## 4 Discussions and Conclusion

It is well-recognized that large visual-language models are important, especially so in the medical imaging field. As mentioned in the introduction, the development of such models to take high-dimensional tomographic images is challenging. This study is the first in this direction, which only used public CT scans and reports and should be easy to reproduce. To improve the performance and validate the generalizability, more big datasets are needed, and the barriers need to be overcome so that patient text reports and genomic data can be shared under IRB approvals. In this regard, follow-up results will be reported in the course of this project, and we are open to more collaboration.

Open-source practice is important for the development of large models. While OpenAI and Google systems are not open sources, recently Cerebras published seven GPT models under the Apache 2.0 license, which have from 111M to 13B parameters and are available on Hugging Face and in Model Zoo on GitHub. Also, a scaling law was derived using these models on the Pile dataset linking compute to accuracy. The Pile is an 800GB dataset of diverse text files, an ideal benchmark for natural language processing. Up to now, Cerebras-GPT is the only industry-leading family of open-source models. Forbes wrote on March 28, 2023, that “By releasing these seven GPT models, Cerebras not only demonstrates the power of its CS-2 Andromeda supercomputer as being amongst the premier training platforms but elevates Cerebras researchers to the upper echelon of AI practitioners.” After our large models are tested at scale and developed into a user-friendly form, we will publicly share our implementation details and codes.

In conclusion, we have performed a feasibility study on building a multi-task CT learning model for lung cancer diagnosis by combining cone-beam CT scans with radiology reports in the domain of public data. Our model consists of LLM and LIM models to perceive multi-modal information under task-specific text prompts for optimized diagnostic performance. Our initial results show that the proposed model performs multiple medical tasks well, including lung segmentation, lung nodule detection, and lung cancer classification. Active efforts are in progress.

## Footnotes

1 https://cdas.cancer.gov/nlst/

2 https://doi.org/10.7937/TCIA.2020.NNC2-0461

